# Molecular dissection of the structural and nonstructural proteins of spanish-1918 Influenza, pandemic-2009, and bird flu viruses

**DOI:** 10.1101/585778

**Authors:** Gusti Ngurah Mahardika, Nyoman Suartha, Gusti Ayu Yuniati Kencana, Ida Bagus Kade Suardana, Nyoman Sri Budayanti

**Author notes:** Corresponding Author: Gusti Ngurah Mahardika, Virology Laboratory, Faculty of Veterinary Medicine, Udayana University, Jl. PB Sudirman, Denpasar, Bali, Indonesia.

## Abstract

The potential emergence of deadly pandemic influenza viruses is unpredictable and most have emerged with no forewarning. The distinct epidemiological and pathological patterns of the Spanish (H1N1), pandemic-2009 (H1N1), and avian influenza (H5N1), known as bird flu, viruses may allow us to develop a ‘template’ for possible emergence of devastating pandemic strains. Here, we provide a detailed molecular dissection of the structural and nonstructural proteins of this triad of viruses. GenBank data for three representative strains were analyzed to determine the polymorphic amino acids, genetic distances, and isoelectric points, hydrophobicity plot, and protein modeling of various proteins. We propose that the most devastating pandemic strains may have full-length PB1-F2 protein with unique residues, highly cleavable HA, and a basic NS1. Any newly emerging strain should be compared with these three strains, so that resources can be directed appropriately.

## Background

We must be constantly alert to the potential emergence of deadly pandemic influenza viruses because most have emerged with no forewarning. The emergence of new strains will continue to challenge public health and pose problems for scientific communities ^1^. The distinct epidemiological patterns of the Spanish (H1N1), pandemic-2009 (H1N1), and bird flu (H5N1) viruses should allow us to develop a ‘template’ for possibly devastating pandemic strains. Spanish flu and pandemic-2009 flu established human-to-human infections ^2,3^, whereas bird flu, the highly pathogenic avian influenza virus (HPAIV) subtype H5N1 (HPAIV-H5N1), has so far shown only limited or unsustainable human-to-human transmission ^4–6^. These three strains have spread worldwide, although bird flu has yet to arrive on the American continent ^7^, while this is still theoretically possible ^8^.

Spanish flu, which caused an estimated 20–50 million deaths around 1918 ^9^, is recognized as the most devastating epidemic in modern history ^10^. However, the case fatality rate (CFR) of Spanish flu is not the highest recorded, although its global CFR is believed to have been around 2.5% ^3^. The CFR of pandemic-2009 is estimated to have been approximately 0.05% of confirmed cases, so its virulence is deemed mild ^11^. The CFR increases greatly if the denominator is influenza-related critical illness. The mortality rate associated with critical illness in pandemic-2009 in various countries was reported to range from 15% in Australia to 61% in Southeast Asia ^12^. Among the viral triad, the CFR of bird flu is highest. As of December 19, 2016, the cumulative number of confirmed human cases of avian influenza A (H5N1) reported to the World Health Organization in 2003–2006 was 856, 452 (52.8%) of which were fatal (www.who.int). However, this comparison of mortality rates is invalid because the availability of and access to medical interventions were significantly different in 1918 from now. The fatalities associated with pandemic-2009 and bird flu may have been much higher without modern medical modalities.

The influenza viruses are a unique family of viruses, the *Orthomyxoviridae*, and members have an eight-segment negative-strand genome ^13^. Five segments encode single proteins: (from the largest to smallest) polymerase basic 2 (PB2), polymerase acidic (PA), hemagglutinin (HA) nucleoprotein (NP), and neuraminidase (NA) ^13^. The second largest fragment encodes polymerase basic 1 (PB1) ^13^ and an accessory peptide, PB1-F2 ^14,15^. The two smallest segments are each spliced to produce the mRNAs for two proteins, M1 and M2 (the seventh segment) and NS1 and NS2 (the eighth segment) ^13^.

No direct molecular comparison of all the genes of this influenza virus triad has been made. Molecular modeling of NA has been reported, which established the susceptibility patterns of the viruses to anti-influenza drugs ^16^. A phylogenetic analysis and amino acid homology analysis of the polymerase complex genes of Spanish flu, seasonal flu, and bird flu have been described, and concluded that the genes of Spanish Flu closely resemble those of bird flu ^17^. Although the analysis of viral genomes alone is unlikely to clarify some critical issues ^3^, such as the viral capacity for human-to-human transmission and the severity of clinical infection, a head-to-head molecular comparison of different lineages should pinpoint the putative gene(s) or protein(s) that can be used as signals for the potential emergence of a catastrophic pandemic strain. Here, we provide a detailed molecular analysis of the structural and nonstructural proteins of this triad of viruses that are associated with their distinct transmissibility and severity.

## Results

The genetic distances, numbers of amino acid differences, and isoelectric points of all the structural and nonstructural proteins of the SF, PDM, and BF viruses are presented in Table 2. The lengths of all the corresponding proteins of the triad were equal, except those of PB1-F2, NS1, and HA. In SF and BF, PB1-F2 contains 90 residues, whereas in PDM, it contains only 11 residues. NS1 in PDM has lost 11 amino acids at the carboxyl terminus. The length of HA in SF and PDM is 566 amino acids, whereas in BF, it is 568 amino acids. The genetic distance between BF and SF is smaller than that between PDM and SF based on PB1 and PA, whereas the contrary true when the distances are based on PB2, HA, NS1, or NS2. The genetic distances are almost equal between BF and SF and between PDM and SF when based on other protein-coding fragments. Fewer amino acid differences are in all protein of BF to SF than PDM to SF, except in HA, NS1, and NS2. The isoelectric points of all proteins are almost equal in the three viruses, except that HA of PDM and NS1 of SF are more basic than the corresponding proteins in the other viruses.

The superimposed hydrophobicity plots of all the proteins are given in Supplementary Material 1. The plots of PB1-F2, HA, NS1, and NS2 of SF, PDM, and BF are presented in Figure 1. The plots for PB2, PB1, PA, NP, NA, MA1, and MA2 were almost perfectly superimposed. A slight aberration was observed in HA, which is neutral in SF and PDM, but hydrophilic in BF at residues 300–400 (Figure 1). PB1-F2 is also more hydrophilic in BF than in SF at positions 1–20 and 60–70, but more hydrophobic at positions 30–50. The NS1 and NS2 plots were not superimposed exactly, and the mismatch was most prominent in the BF plot (Figure 1).

**Figure 1.**
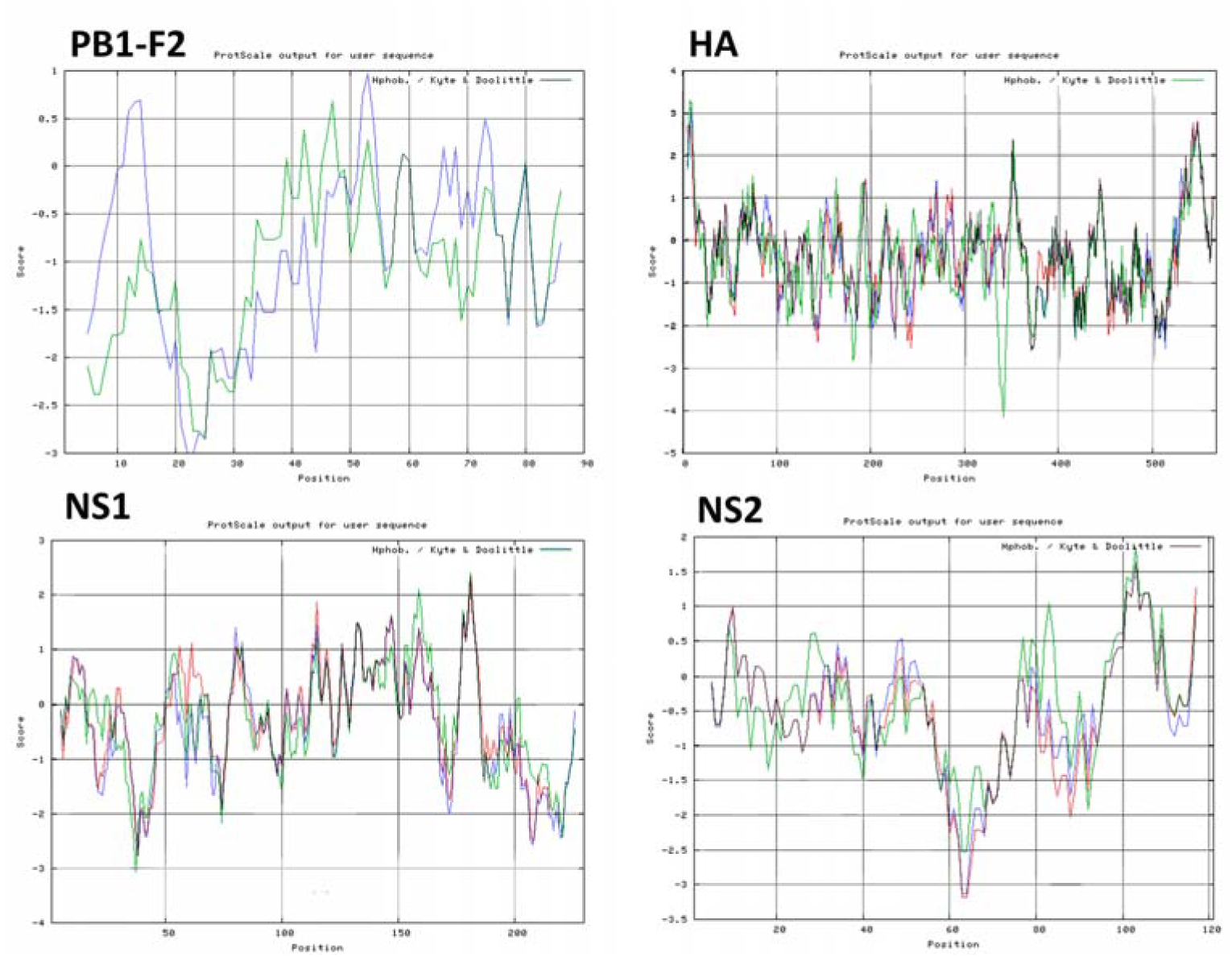
Superimposed hydrophobicity plots of Spanish (blue), pandemic-2009 (red), and bird flu (green) viruses for PB1-F2 (top left), HA (top right), NS1 (bottom left), and NS2 (bottom right). Hydrophobicity plot of each peptide was calculated with the Kyte and Doolittle method ^44^ using the online software ProtScale ^45^. (http://web.expasy.org/protscale/), and the results were superimposed with Adobe Creative Cloud 2015 (Adobe System Inc.).

The result of protein modeling of PB1-F2, NS1, and NS2 are presented in Figures 2–4. PB1-F2 and NS2 are dominated by an α-helix, with no β-sheet. NS1 consists of an α-helix membrane domain and a globular head consisting of an α-helix and a β-sheet. All those viral proteins have homologous structures, with some minor differences. PB1-F2 consists of two α-helixes. The amino-terminal helix is continuous in BF, whereas it is interrupted in SF. On the contrary, the carboxyl-terminal helix of SF is continuous, whereas that of BF is interrupted by random coils. NS1 consists of a globular head formed by the amino terminus of the protein, and a trans-membrane domain at the carboxyl terminus. NS2 consists of four α-helix structures. The first helix (from the amino terminus) is intact in BF, but interrupted in SF and PDM. The second (yellow in Figure 4) is intact in SF and BF, but interrupted in PDM. The third (green to light blue in Figure 4) is intact in all three viruses. The last helix at the carboxyl terminus (blue in Figure 4) is intact in SF, but interrupted in PDM and BF.

**Figure 2.**
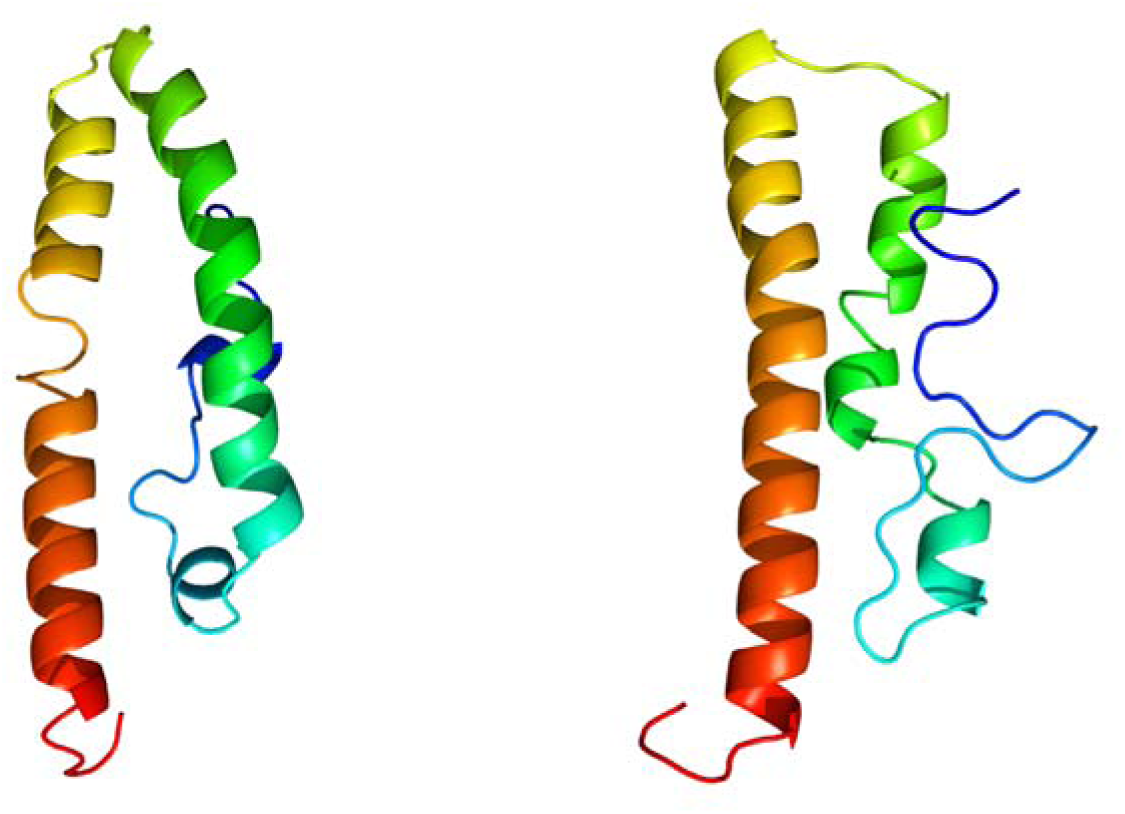
Cartoon peptide modeling of PB1-F2 of SF (left) and BF (right). Images are colored by inverted rainbow from N- to C-terminus. Protein modeling was performed with the online resource PYRE2 (http://www.sbg.bio.ic.ac.uk) ^46^. Protein models were visualized with RasWin 2.7.5.2 (www.rasmol.org).

**Figure 3.**
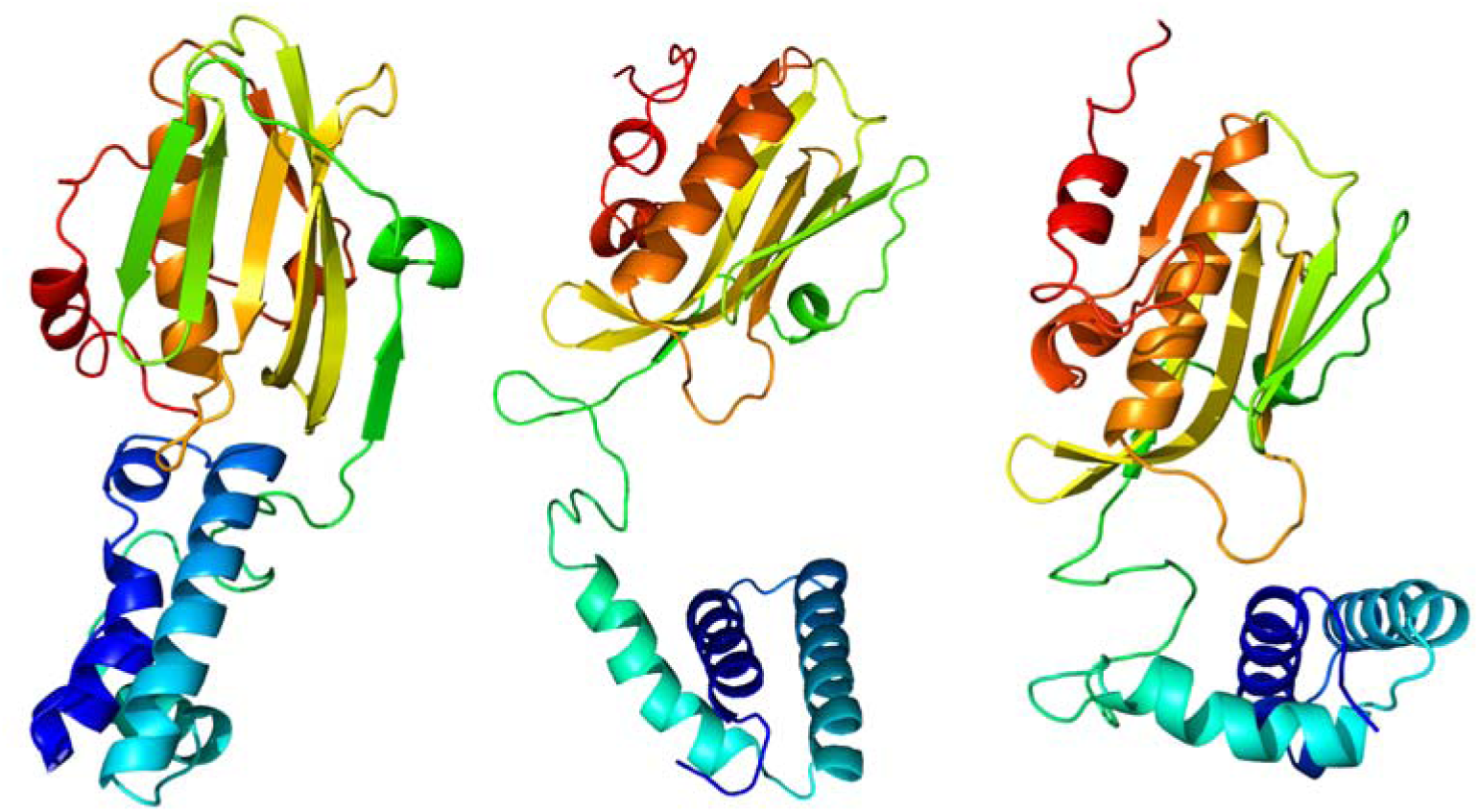
Cartoon peptide modeling of NS1 of SF (left), PDM (middle), and BF (right). Images are colored by inverted rainbow from N- to C-terminus. Protein modeling was performed with the online resource PYRE2 (http://www.sbg.bio.ic.ac.uk) ^46^. Protein models were visualized with RasWin 2.7.5.2 (www.rasmol.org).

**Figure 4.**
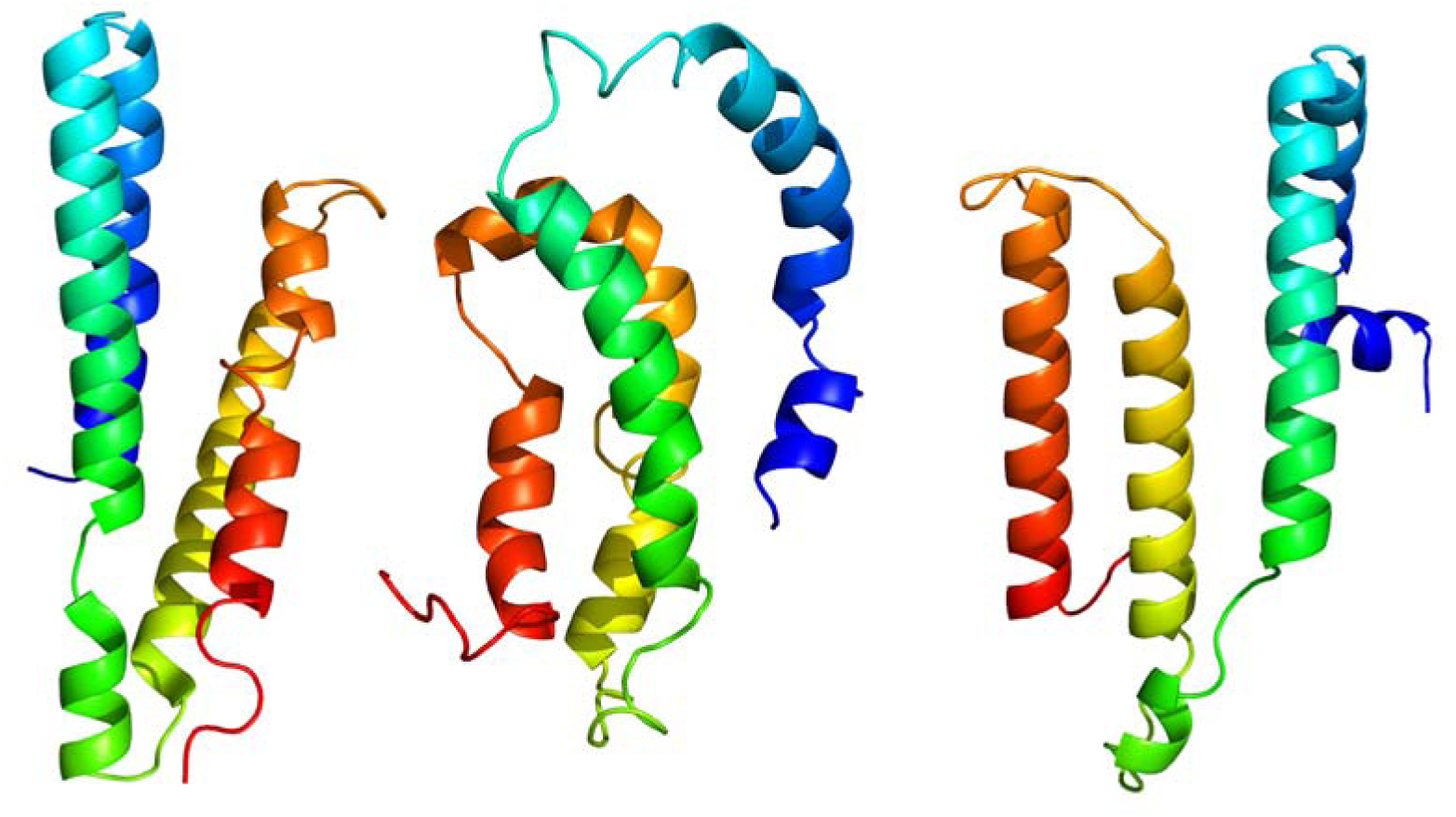
Cartoon peptide modeling of NS2 of SF (left), PDM (middle), and BF (right). Images are colored by inverted rainbow from N- to C-terminus. Protein modeling was performed with the online resource PYRE2 (http://www.sbg.bio.ic.ac.uk) ^46^. Protein models were visualized with RasWin 2.7.5.2 (www.rasmol.org).

## Discussion

It is generally believed that the pathogenicity and transmissibility of influenza viruses are polygenic or multifactorial ^18,19^. Many gene segments of these viruses contribute to their capacity to cause severe outcomes in their host and to be readily transmitted between hosts ^20^. Whereas one viral protein might define a pathogenic characteristic, virulent outcome may only be possible with the contribution of other proteins. The magnitude of an infection depends on those proteins acting in concert with host and environmental factors ^21–24^. In other word, the genetic make-up of the virus must match the permissiveness of the host and an appropriate environment to generate fatal or contagious outcomes.

The full genomic sequence of the Spanish-1918 influenza virus, which was derived from a naturally preserved human body believed to have died in that severe pandemic ^17,25^, and the massive number of influenza virus sequences determined in recent decades provide data from which a ‘template’ for the emergence of modern disastrous pandemic strains can be generated.

The triad strains SF, PDM, and BF are representative of different influenza viruses with distinct hallmarks. All have been disseminated globally, except BF, which so far affects only Asia, Europe, and Africa ^7^. The SF and PDM viruses are readily transmitted between human, whereas BF has no capacity for sustained human-to-human transmission ^4–6^. BF has the highest CFR, at >50% (www.who.int). Without modern medical resources, as occurred in 1918, its CFR might be much higher. Spanish-1918 influenza, responsible for the ‘mother of pandemics’, had a CFR of 2%–3% ^3^. Compared with those viruses, the pandemic-2009 virus is ‘avirulent’ or mild ^11^, with a CFR of only 0.05% ^11^.

We analyzed the genetic distances between the triad viruses, the differences in the numbers of amino acids and the isoelectric points of all their structural and nonstructural proteins, as well as their hydrophobicity plots. We found that BF closely resembles SF, except in proteins HA and NS2. In those proteins, PDM is closer to SF than is BF. We identified PB1-F2, HA, and NS1 are the factors putatively responsible for the distinct human-to-human transmissibility of SF, PDM, and BF, and as hallmarks of disease severity.

PB1-F2 seems to be responsible for pathogenicity. PB1-F2 contains 90 residues in SF and BF, but only 11 residues in PDM. Therefore, the mild disease associated with PDM might be related to the loss of PB1-F2. Different hydrophobicity patterns might also be responsible for the differences between SF and BF. PB1-F2 is more hydrophilic at positions 1–20 and 60–70 in BF than in SF, but more hydrophobic at positions 30–50.

The obvious differences in the open reading frame (ORF) length of this peptide in SF, PDM, and BF suggest that PB1-F2 is responsible for the pathological outcomes of influenza viral infections. The truncation of PB1-F2 in PDM might have caused the low CFR of PDM in 2009. PB1-F2 is a small accessory protein encoded by an alternative ORF in the second largest segment of most influenza A virus genomes ^26^ and is thought to contribute to viral pathogenicity and the severity of pandemic influenza ^15^.

The role of PB1-F2 in the pathogenesis of the influenza viruses is contentious. The function of this peptide is both strain- and host-specific ^27^, and it is even expressed strain-specifically ^28^. In various studies, this protein caused immune disruption through the recruitment of neutrophils and the inhibition of natural killer cells ^29^, immune cell apoptosis ^26^, and the inhibition of interferon ^30^. Its species-specific activity was demonstrated in the attenuation of AIV-H5N1 ^31^ and in an experiment with swine isolates ^28^. The posttranslational phosphorylation of Ser^35^ ^32^ and Ser^66^ ^33^ of PB1-F2 may contribute to the strain-specific functions this protein. Both SF and BF have serine at residue 35, whereas Ser^66^ in SF is substituted with asparagine in BF.

The hydrophilic domain around the cleavage site of HA may be responsible for the highly pathogenic nature of BF. HA contains 566 amino acids in SF and PDM, but 568 amino acids in BF. The superimposition of the hydrophobicity plots of HA showed slight aberrations, with more hydrophilic amino acids in the region defined by residues 300–400 in BF than in the other viruses. This domain includes the cleavage site of HA.

HA is an important surface protein for receptor recognition and the penetration of the virus to the host cell cytoplasm ^13^. Cleavage of this virus is a prerequisite for viral infections and is therefore a crucial determinant in viral pathogenicity and tissue tropism ^34^. The most highly pathogenic strains have polybasic amino acids at the cleavage site or no carbohydrate residues in the vicinity of the site ^18^. The length of the site also enhances its cleavability ^18^. The long track of basic amino acids at the cleavage site of HA in HPAIVs is thought to facilitate the expansion of tissue tropism. The cleavage of the HA precursor to HA1 and HA2 is mediated by a ubiquitous endopeptidase, furin, located in the trans-Golgi network ^19^. Infectious virions are released from infected cells with no requirement for extracellular activation. Moreover, the cleavability and pathogenicity of HA seem to be strain- and host-specific. Zhang et al. showed experimentally that a single substitution in the cleavage site of HA modulates the virulence of the H5N1 virus ^35^, and another report ^36^ demonstrated that the contribution of the H5 multibasic site to the virulence of the HPAIV-H5N1 virus varies among mammalian hosts, and is most significant in mice and ferrets and less remarkable in nonhuman primates. The postulate of Horimoto and Kawaoka ^37^ that the cleavability of HA correlates with the degree of virulence, when all other genetic characteristics are considered equal, is valid. However, in the evolution of a low-pathogenic avian influenza virus to become an HPAIV, a change in the cleavage site alone is not enough. The low-pathogenic strains may already have cryptic virulence potential ^38^.

NS1 and NS2 may be responsible for the human-to-human transmission of the influenza virus. The PDM virus resembles SF more closely than BF. The hydrophobicity plots of NS1 and NS2 were incompletely superimposed, and the aberration was predominantly located in the BF plot (Figure 1). NS1 and NS2 are encoded by the shortest segment of the influenza genome. NS1 is a nonstructural protein expressed in high abundance in infected cells, whereas NS2 seems to be a structural protein that is a minor component of the virion ^13^. NS1 is collinearly translated from the transcript of the segment, whereas NS2 is encoded by a spliced transcript ^13^. NS1 is particularly linked to the ‘cytokine storm’ phenomenon that follows influenza infection ^39^, contributing to its unusual immunopathogenesis ^40^. The hyperinduction of proinflammatory cytokines is a hallmark of severe influenza infection ^41^. The presence of host cofactors determines the effect of this protein. NS1 only induces a cytokine storm in the presence of the myeloperoxidase system ^39^, but the mechanism of NS2 still requires clarification. NS1 of SF is remarkably basic, whereas NS1 of the other viruses is acidic (Table 2). This unique property of SF NS1 might have led to the devastating humanitarian impact of the SF pandemic shortly before the 1920s.

Amino acid differences disturb of secondary structures of PB1, NS1, and NS2, which might alter the human-to-human transmissibility and human pathogenicity of this triad of viruses. The modeling of the PB1-F2, NS1, and NS2 proteins showed that the proteins of all three viruses have homologous structures, with some minor differences.

The findings of this study are merely hypothetical. Reverse genetic experiments with combined gene segments are the only way to validate our hypotheses. Such experiments will be controversial and should be strictly regulated. A wide survey of the influenza virus genomes available in databases will offer indirect evidence to support our findings. The analysis conducted in this study was very simple and could be undertaken in many countries, so a capacity to immediately predict the potential impact of an emergent strain is possible in these countries.

We conclude that the putative pathogenicity of an influenza virus lies in PB1-F2 and the cleavability of HA, whereas NS1 and NS2 (especially NS1) are responsible for the human permissiveness of the virus. The most devastating pandemic strains may have full-length PB1-F2 proteins with unique residues, highly cleavable HA, and a basic NS1. The generation of such strain with reverse genetics will provide proof of this model. However, this kind of experiment must be strictly regulated or may even be impossible. Any newly emerging strain should be compared with this triad of influenza viruses to rapidly estimate its pathogenicity and human-to-human transmissibility.

## Materials and Methods

The nucleotide and deduced amino acid sequences of all the structural and nonstructural proteins (PB2, PB1, PB1-F2, HA, NP, NA, M1, M2, NS1, and NS2) of Spanish Flu, pandemic-2009, and bird flu were analyzed in this study. Strain A/Brevig Mission/1/1918(H1N1) provided all the genomic segments of Spanish flu, except the HA gene, and strain A/South Carolina/1/18 (H1N1) provided the HA gene. The A/California/04-061-MA/2009(H1N1) strain represented the pandemic-2009 virus, and strain A/goose/Guangdong/1/1996(H5N1) represented the bird flu virus. The viruses are abbreviated hereinafter to SF, PDM, and BF for Spanish flu, pandemic-2009, and bird flu, respectively. The accession numbers for the GenBank data are presented in Table 1. The sequences were aligned using ClustalW, and the polymorphic amino acids in each protein and the numbers of amino acid differences were determined with Mega 6.0 ^42^. Genetic distances were calculated with the Kimura 2-parameter model ^43^. The isoelectric points were calculated with Peptide Calculator (http://www.bachem.com). A hydrophobicity plots of each peptide was constructed with the Kyte and Doolittle method ^44^, using the online software ProtScale ^45^ (http://web.expasy.org/protscale/), and the results were superimposed using Adobe Creative Cloud 2015 (Adobe System Incorporated). Protein modeling was performed with the online resource PYRE2 (http://www.sbg.bio.ic.ac.uk) ^46^, and the protein models were visualized with RasWin 2.7.5.2 (www.rasmol.org).

**Table 1.**
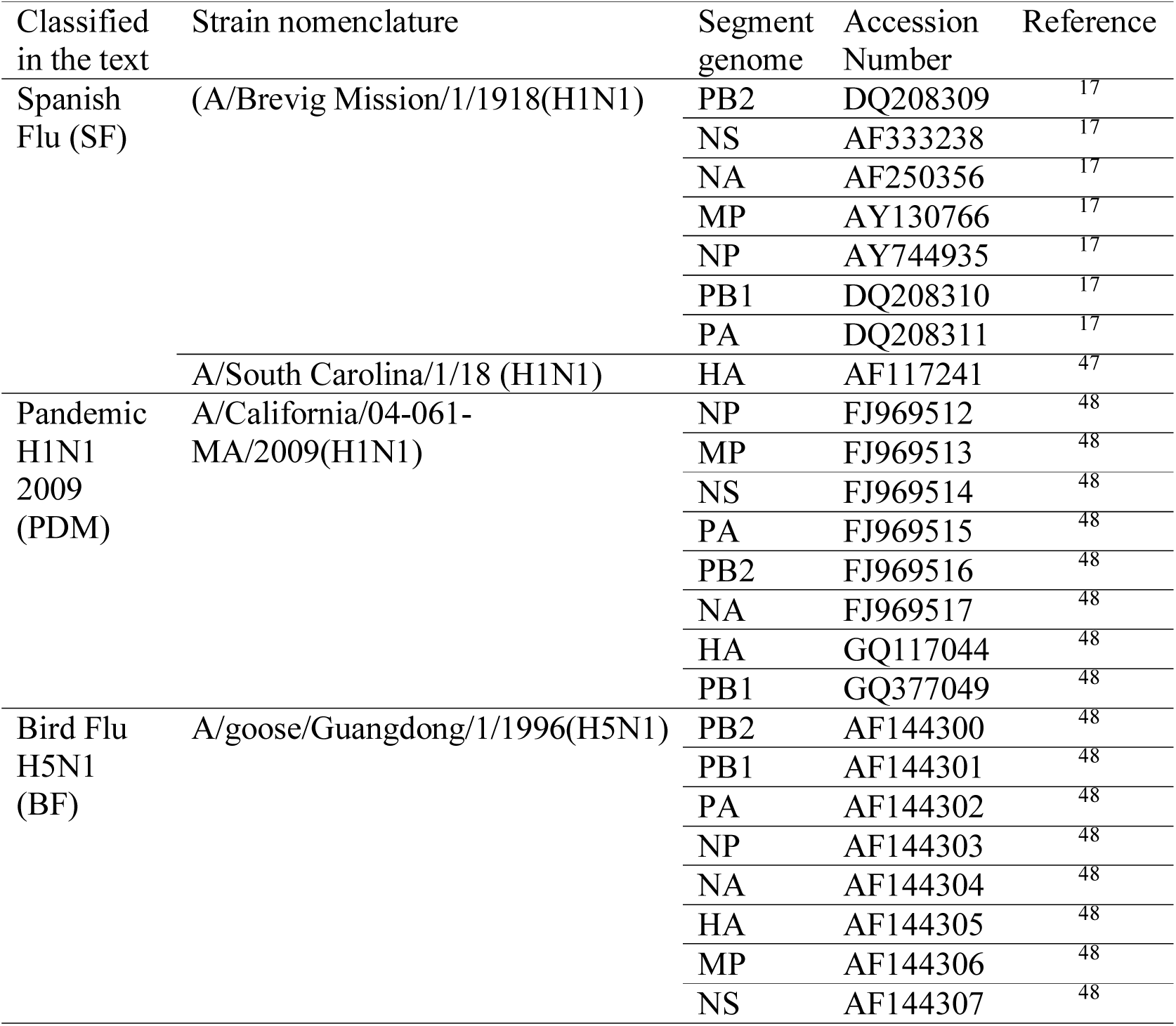
GenBank sequence data for Spanish flu, pandemic-2009 H1N1, and bird flu H5N1 analyzed in this study

**Table 2.** Genetic distances, numbers of amino acid differences, and isoelectric points of all structural and nonstructural proteins of Spanish-H1N1, pandemic-2009 H1N1, and bird flu-H5N1 viruses.

| Gene | Protein length<br>(residues) |  |  | Genetic distance |  | Number of amino<br>acid difference |  | Isoelectric point |  |  |
| --- | --- | --- | --- | --- | --- | --- | --- | --- | --- | --- |
|  | SF | PDM | BF | SF-PDM | SF-BF | SF-PDM | SF-BF | SF | PDM | BF |
| PB2 | 759 | 759 | 759 | 0.14 | 0.18 | 25 | 19 | 9.9 | 9.8 | 9.7 |
| PB1 | 757 | 757 | 757 | 0.21 | 0.18 | 28 | 19 | 9.5 | 9.6 | 9.5 |
| PB1-F2 | 90 | 11 | 90 | NA | 0.11 | 27 | NA | 10.7 | NA | 11 |
| PA | 716 | 716 | 716 | 0.18 | 0.16 | 25 | 24 | 5.3 | 5.3 | 5.4 |
| HA | 566 | 566 | 568 | 0.22 | 0.49 | 80 | 195 | 6 | 6.9 | 6.3 |
| NP | 498 | 498 | 498 | 0.13 | 0.14 | 28 | 10 | 9.5 | 9.6 | 9.7 |
| NA | 469 | 469 | 469 | 0.2 | 0.19 | 59 | 44 | 5.4 | 6.1 | 5.9 |
| M1 | 252 | 252 | 252 | 0.11 | 0.1 | 11 | 5 | 9.6 | 9.8 | 9.8 |
| M2 | 97 | 97 | 97 | 0.06 | 0.06 | 8 | 6 | 4.9 | 4.5 | 4.6 |
| NS1 | 230 | 219 | 230 | 0.17 | 0.4 | 37 | 72 | 8.2 | 5.7 | 6.4 |
| NS2 | 121 | 121 | 121 | 0.12 | 0.21 | 12 | 20 | 5 | 5.1 | 4.8 |

Genetic distances were calculated with the Kimura 2-parameter model <sup>49</sup>. Isoelectric points were calculated with Peptide Calculator (http://www.bachem.com).
SF: Spanish-H1N1; PDM: pandemic-2009 H1N1; BF: bird flu-H5N1.

## Acknowledgments

We thank Mr. Wisnu Wira Mahardika, a student at Bina Nusantara University in Tanggerang–Banten, Indonesia, for assistance in generating the computer graphics. All authors are member of research team funded by Professor Publication and Promotion Project Udayana University of Bali, DIPA-PNBP 2018, Grant No. **383-1/UN14.4.A/LT/2018** with GNM as Principal Investigator. GAYK, IBKS and GNM are member of research team funded by The National Key Research and Development Program of China, Grant No. 2016YFE0203200; in which GNM is Indonesian Principal Investigator. The funders have no role in drafting and finalizing the document and in making decision to publish.

## Additional information

### Compliance with Ethics Guidelines

This article does not contain any studies with human participants or animals performed by any of the authors.

## Author contributions

GNM and NSB conceived of or designed study. NSB, NS, GAYK and IBKS performed data collection. All author analyzed data. GNM and NSB wrote the paper.

## Supplementary figure

Superimposed hydrophobicity plots of Spanish (blue), pandemic-2009 (red), and bird flu (green) viruses for all proteins. Hydrophobicity plot of each peptide was calculated with the Kyte and Doolittle method ^44^ using the online software ProtScale ^45^ (http://web.expasy.org/protscale/), and the results were superimposed with Adobe Creative Cloud 2015 (Adobe System Inc.). First row: PB2 (left), PB1 (middle), PB1-F2 (right); second row: PA (left), HA (middle), NP (right); third row: NA (left), M1 (middle), M2 (right); forth row: NS1 (left), NS2 (middle).

