## Supplementary figures and images for "Molecular dissection of the structural and nonstructural proteins of spanish-1918 Influenza, pandemic-2009, and bird flu viruses"

### Supplementary figure

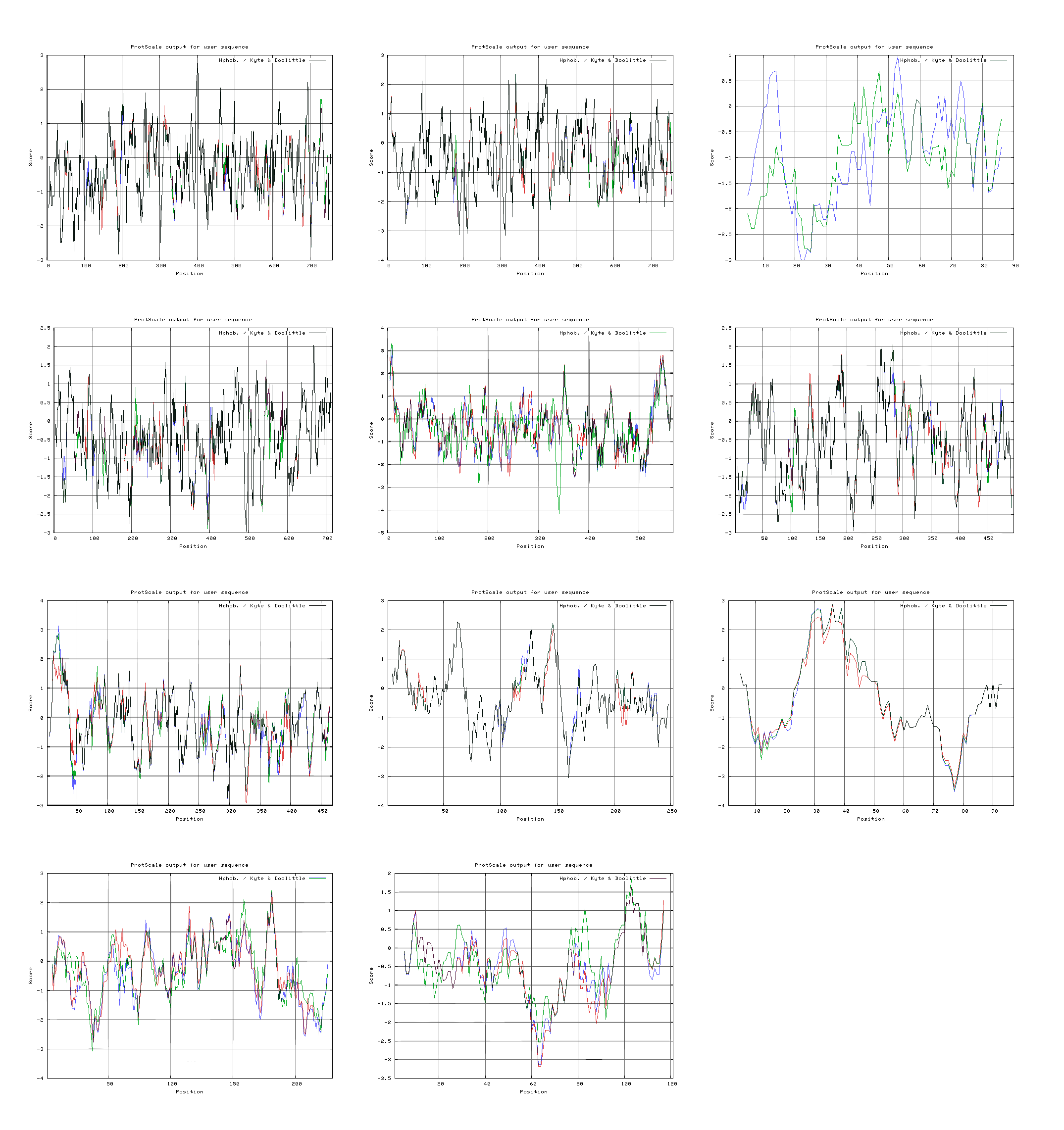
